# Bioenergetic function is decreased in peripheral blood mononuclear cells of veterans with Gulf War Illness

**DOI:** 10.1101/2023.06.07.544068

**Authors:** Joel N Meyer, William K Pan, Ian T Ryde, Thomas Alexander, Jacquelyn C. Klein-Adams, Duncan S. Ndirangu, Michael J Falvo

**Affiliations:** Nicholas School of the Environment, Duke University, Durham, NC; Department of Veterans Affairs, War Related Illness and Injury Study Center, East Orange, NJ; New Jersey Medical School, Rutgers Biomedical and Health Sciences, Newark, NJ

## Abstract

Gulf War Illness (GWI) is a major health problem for approximately 250,000 Gulf War (GW) veterans, but the etiology of GWI is unclear. We hypothesized that mitochondrial dysfunction is an important contributor to GWI, based on the similarity of some GWI symptoms to those occurring in some mitochondrial diseases; the plausibility that certain pollutants to which GW veterans were exposed affect mitochondria; mitochondrial effects observed in studies in laboratory models of GWI; and previous evidence of mitochondrial outcomes in studies in GW veterans. A primary role of mitochondria is generation of energy via oxidative phosphorylation. However, direct assessment of mitochondrial respiration, reflecting oxidative phosphorylation, has not been carried out in veterans with GWI. In this case-control observational study, we tested multiple measures of mitochondrial function and integrity in a cohort of 114 GW veterans, 80 with and 34 without GWI as assessed by the Kansas definition. In circulating white blood cells, we analyzed multiple measures of mitochondrial respiration and extracellular acidification, a proxy for non-aerobic energy generation; mitochondrial DNA (mtDNA) copy number; mtDNA damage; and nuclear DNA damage. We also collected detailed survey data on demographics; deployment; self-reported exposure to pesticides, pyridostigmine bromide, and chemical and biological warfare agents; and current biometrics, health and activity levels. We observed a 9% increase in mtDNA content in blood in veterans with GWI, but did not detect differences in DNA damage. Basal and ATP-linked oxygen consumption were respectively 42% and 47% higher in veterans without GWI, after adjustment for mtDNA amount. We did not find evidence for a compensatory increase in anaerobic energy generation: extracellular acidification was also lower in GWI (12% lower at baseline). A subset of 27 and 26 veterans returned for second and third visits, allowing us to measure stability of mitochondrial parameters over time. mtDNA CN, mtDNA damage, ATP-linked OCR, and spare respiratory capacity were moderately replicable over time, with intraclass correlation coefficients of 0.43, 0.44, 0.50, and 0.57, respectively. Other measures showed higher visit-to-visit variability. Many measurements showed lower replicability over time among veterans with GWI compared to veterans without GWI. Finally, we found a strong association between recalled exposure to pesticides, pyridostigmine bromide, and chemical and biological warfare agents and GWI (p < 0.01, p < 0.01, and p < 0.0001, respectively). Our results demonstrate decreased mitochondrial respiratory function as well as decreased glycolytic activity, both of which are consistent with decreased energy availability, in peripheral blood mononuclear cells in veterans with GWI.

## Introduction

Gulf War Illness (GWI) is a major health problem for approximately 250,000 Gulf War veterans (GVs) from the United States, comprising roughly one-third of all US veterans who served in the Persian Gulf War [1]. There is also evidence that veterans from other countries were affected [2, 3]. However, the etiology of GWI is unclear. The clinical presentation of GWI is heterogeneous, but is characterized across multiple organ systems with common chronic symptoms including exercise intolerance, fatigue, pain, and neurocognitive symptoms. Three theoretical considerations suggest that mitochondrial dysfunction is a driver of GWI symptoms. First, many GWI symptoms are consistent with multisystem symptoms displaying clinical heterogeneity and tissue- specific manifestations observed in civilians with genetically-based mitochondrial disorders [4]. Second, many of the tissues affected in GWI are high energy use (e.g., muscular, central/autonomic nervous, respiratory) [5, 6]. This is consistent with the hypothesis that many symptoms result from mitochondrial dysfunction, due to the essential role of mitochondria in energy production: a primary role of mitochondria is production of energy in the form of ATP, via oxidative phosphorylation (**Fig. 1**). Tissues and organs that rely predominantly on mitochondrial oxidative phosphorylation for energy production are often those that exhibit the greatest pathology when mitochondrial function is compromised [7]. Third, mitochondria are a sensitive target of many chemicals [8–10], including many of those to which GVs were exposed. These include widely-used pesticides such as permethrin [11–13], chlorpyrifos [14–17], malathion [18–20]; polycyclic aromatic hydrocarbons resulting from incomplete combustion, which cause high levels of damage to the mitochondrial genome (mtDNA) [21, 22] that is not repaired due to the absence of the relevant DNA repair pathway [23, 24]; particulate matter [25–27]; depleted uranium [28]; and possibly pyridostigmine bromide (PB, a carbamate drug intended to protect against nerve gas attacks), especially in combination with permethrin and other relevant exposures [29–32]. Several studies that employ combinations of some of these chemicals to create rodent models of GWI also reported alterations in mitochondria function [31, 33, 34].

**Figure 1.**
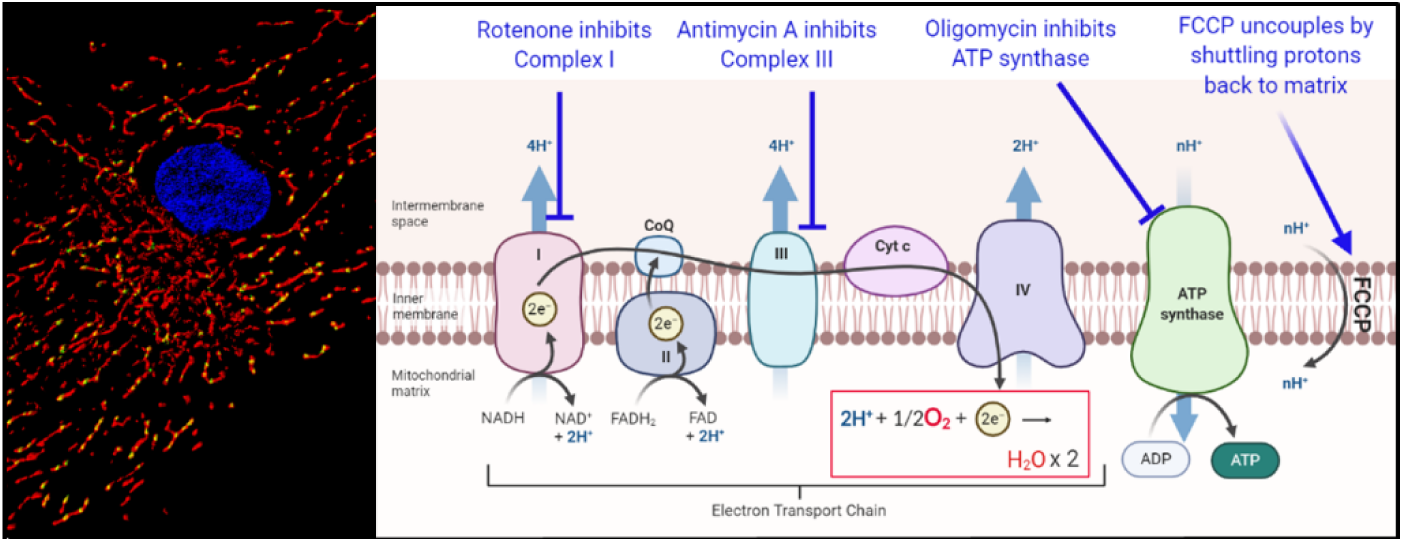
Schematic illustrating mitochondrial parameters analyzed: mtDNA copy number (CN). mtDNA damage, mitochondrial and non-mitochondrial oxygen consumption. Left panel from Meyer et al·, 2013^6^ shows the many mtPNAs in a cell in yellow (mitochondria in red, nucleus in blue). The number of mtDNAs is stress-responsive. Decreased number or damage to mtDNA could interfere with proper assembly of the electron transport chain (right panel), because components of Complexes I, III, IV, and ATP synthase are encoded in mtDNA; increased mtDNA CN may indicate a stress-responsive attempt to compensate for decreased function. The electron transport chain converts food-derived fuel to a proton gradient by pumping protons out of the matrix. ATP synthase uses this proton gradient to generate energy in the form of ATP. Complex IV converts oxygen to water (highlighted and boxed in red); thus, measurement of oxygen consumption is a powerful measure of mitochondrial function- mediated oxygen consumption. The amount of oxygen used to make ATP is measured by inhibiting ATP synthase with oligomycin. The maximal amount of oxygen that mitochondria can consume when function increases is measured by “uncoupling” with FCCP; FCCP shuttles protons back to the matrix without any production of ATP, thus uncoupling oxygen consumption from ATP production. Finally, the contribution of non-mitochondrial processes to oxygen consumption is measured by addition of rotenone and ant«mycin A, which inhibit all electron flow and therefore all mitochondrial oxygen consumption. Any oxygen consumption that occurs in the presence of oligomycin that is mitochondrial can be attributed to proton leak.

In addition to these theoretical considerations, multiple publications now demonstrate that a variety of indicators of mitochondrial function are altered in blood samples from veterans with GWI. Measures have included blood lactate [35] and calf muscle phosphocreatine [36] post-exercise; mtDNA damage [37]; changes to the amount of mtDNA per cell [37]; electron transport chain Complex I activity [37]; and metabolomic alterations consistent with decreased mitochondrial beta-oxidation [38]. In addition to these studies in GVs, support has also come from rodent models of GWI, in which researchers have identified decreased mitochondrial lipids (long-chain acyl carnitines), disrupted mitochondrial proteomics, and alterations in transcriptomic patterns indicative of inflammation, oxidative stress, and mitochondrial dysfunction [31, 33, 34]. Finally, publications have demonstrated that a mitochondrial- targeted antioxidant [39] and metabolic activator [40] were protective, which is consistent with a role for dysfunctional mitochondria in GWI. This literature was recently reviewed [41].

Together, these results support the involvement of mitochondria and bioenergetic function in GWI. However, previous studies in GVs have mostly relied on indirect reporters of mitochondrial function, and/or been limited in size (number of GVs tested). To improve our understanding of bioenergetic function in GWI, we compared multiple measurements of mitochondrial oxygen consumption rate (OCR) in GW veterans with and without GWI. OCR measurements provide a direct measurement of oxidative phosphorylation (**Fig. 1**, [42–44]), a fundamental aspect of mitochondrial function.

Measurements under different conditions permit assessment of the overall ability of mitochondria to provide energy. Specifically, we measured OCR under standard culture conditions (“basal” oxygen consumption), ATP-linked oxygen consumption (the portion of mitochondrial oxygen consumption used to convert ADP to ATP), maximal oxygen consumption (this serves as a measure of the ability of mitochondria to increase oxygen consumption on demand, which is important for stress response; the increase in OCR from the basal level is termed “spare respiratory capacity”), and proton leak (which can be considered as either a protective function or a measure of mitochondrial inefficiency) [44]. We also measured nonmitochondrial OCR and extracellular acidification rate (ECAR), a measure of glycolysis. Non-mitochondrial OCR may be associated with inflammation, and glycolysis is a non-mitochondrial and non-oxygen consuming mechanism for producing energy which is sometimes increased in the context of dysfunctional mitochondria [44].

We also analyzed the number of mtDNAs per cell (mtDNA CN) and mtDNA damage. mtDNA CN can vary with a variety of conditions, including mitochondrial stress and disease; we and others have reviewed its utility as a marker of mitochondrial stress [45, 46]. mtDNA damage can also be an important marker of mitochondrial condition. The mitochondrial genome is located adjacent to the electron transport chain (the major source of reactive oxygen species in most cells), is sensitive to many DNA damaging agents (including indirectly, via toxicant-induced mitochondrial dysfunction-mediated generation of reactive oxygen species), and has fewer repair mechanisms than the nuclear genome; as a result, mtDNA damage can be a useful marker of mitochondrial damage in some cases [8, 9, 24]. Thus, changes in mtDNA CN and mtDNA damage may serve as markers of exposure to a wide array of mitochondrial stressors. Furthermore, while most of the proteins that make up the electron transport chain complexes are encoded in the nuclear genome, 13 are encoded in the mitochondrial genome. Therefore, measurements of oxygen consumption are best interpreted in the context of knowledge of changes in mtDNA CN and damage, because change in mtDNA number or integrity could reduce production of mtDNA-encoded electron transport chain proteins. We made these measurements in whole blood fractions containing a combination of PBMCs and platelets.

We tested the primary hypotheses that mtDNA CN would be decreased, mtDNA damage increased, mitochondrial respiration decreased, and ECAR increased, in GWI. We also tested whether recalled exposure to pesticides, chemical or biological agents, and PB would be associated with GWI. Finally, we carried out statistical analyses testing whether the relationship between these mitochondrial parameters and GWI would be affected by gender, exercise, smoking, alcohol use, and body-mass index, parameters shown in previous work to affect mitochondrial function.

## Materials and Methods

### Participants

Gulf War Veterans (GV) who deployed in support of Operations Desert Storm and Shield (1990-1991) and those who did not deploy but served during this period were recruited between April 2017 and July 2021 to participate in this study through our national specialty clinic (New Jersey War Related Illness and Injury Study Center) and the surrounding region using traditional methods (e.g., research flyers, website advertisement, word of mouth) as well as through rosters provided by the Department of Defense’s Manpower Data Center. GWI case status was determined in accordance with the Kansas GWI Case Definition [47]. In brief, GVs with GWI must endorse moderately severe symptoms in at least three symptom domains (pain, fatigue neurocognitive, skin, gastrointestinal, and/or respiratory), with symptoms first endorsed during or following their Gulf War deployment. GVs who did not meet the GWI case definition served as controls. GVs with or without GWI were excluded if they had current or lifetime bipolar I disorder, psychotic disorders, or mood disorder with psychotic features; evidence of active (or partial remission ≤ 1 year) illicit substance use or fitting criteria for active substance dependence and substance dependence in partial remission; taking medications known to affect mitochondrial parameters (e.g., calcium channel blockers, anti-convulsants, nonsteroidal anti- inflammatory drugs <48 hours, bone marrow suppressants); and/or were diagnosed with chronic conditions (e.g. cancer, autoimmune diseases), metabolic disease (e.g. metabolic syndrome, thyroid dysfunction, cardiovascular, renal, liver), or hematological disorders (e.g., hemoglobinopathy, sickle-cell anemia, malaria, etc.). Where possible, electronic medical records were also used to confirm inclusion and exclusion criteria based upon self-report. All participants provided their written informed consent and study procedures were approved by the Department of Veterans Affairs (VA) New Jersey Health Care System and Duke University Institutional Review Board.

### Participant Characteristics

For each participant, we assessed anthropometric characteristics including height, weight and circumferences of the waist and hip using a Gulick tap measure [48]. Resting blood pressure, heart rate, and pulse oximetry were also obtained in a seated position. Following these assessments, participants completed a series of self-administered surveys using REDCap electronic data capture tools hosted by the Department of Veterans Affairs [49, 50]. Surveys included: i) health and medical history, ii) military deployment history, iii) a modified version of the VA’s Cooperative Studies Program #585 Gulf War Era Cohort and Biorepository questionnaire (inclusive of the Kansas questionnaire), iv) Fatigue Severity Scale [51], and v) International Physical Activity Questionnaire (IPAQ) short-form [52]. Electronic data were assigned a random code that was linked to their personal information and that link was only accessible to the study team at the VA New Jersey Health Care System.

### Bioenergetic profiling

To measure real-time mitochondrial function in GVs, peripherical blood mononuclear cells (PBMCs) were isolated from 50 mLs of blood, drawn from each participant’s arm in the antecubital area by qualified clinical staff using sterile technique for blood processing, and analyzed using the Agilent Seahorse XFp. PBMCs were isolated and frozen overnight as described [53]. Blood samples were assigned the same code used with their corresponding electronic data. In brief, drawn blood was diluted (1:1 with PBS-EDTA (2mM)), layered on top of an equivalent volume of Histopaque 1077 (Sigma-Aldrich), centrifuged at 600 x g for 20 minutes. After completion, autologous plasma samples were taken from the top layer and the “buffy coat” was extracted and washed 3 times with 1x PBS (Sigma-Aldrich) at 300 x g, 200 x g, and 100 x g. The cell pellet was then resuspended in 1.2mL autologous plasma, 0.3ml dimethyl sulfoxide (DMSO, Sigma Aldrich), and 1.5mL of warm Roswell Park Memorial Institute medium (RPMI-1640, ThermoFisher) which was supplemented with 10% fetal bovine serum (FBS, Sigma Aldrich), 2 mM glutamine (Sigma Aldrich), 100 units/ml penicillin (Sigma Aldrich) and 0.1 mg/ml streptomycin (Sigma Aldrich). Final cell suspensions were aliquoted into three 1 ml volumes and frozen overnight in an insulated Mr. Frosty container (Nalgene) containing 250mL of isopropanol alcohol (IPA, Sigma Aldrich). After the 24hr freezing period, cells were thawed and analyzed using the Seahorse XFp using the manufacturer’s recommendations and based on Chacko *et al*. [42]. After thaw, cells were centrifuged at 300 x g for 10 minutes. Cell pellets were then resuspended in 2ml of Seahorse XF Base Medium (Agilent) supplemented with 1mM sodium pyruvate (Sigma Aldrich), 2mM L-glutamine (Sigma Aldrich), and 10 mM glucose (Sigma Aldrich). Samples with viability < 70% and/or showing signs of neutrophile activation (where it was difficult to resuspend) were not analyzed. The cell suspension was diluted to a target density of 300,000 cells per well. Samples were layered into 6 sample wells in the Seahorse XFp culture plate, pre-coated with cell-tac, using a sample volume of 175 µL. Two wells containing only 175 µL of the supplemented Seahorse XF Base Medium were used as background correction wells. After plating, the culture plate was spun at 600 x g for 1 minute to ensure adherence. An even layer of cells was desired and confirmed using a microscope. Wells that did not contain an even layer at the bottom or showed signs that the cells did not adhere to the bottom were not included in the analysis. Seahorse injection ports A, B, and C were filled with 25 µl of 40 µM oligomycin for a final in-well concentration of 5 µM (“Oligo,” VWR Scientific), 4.5 µM carbonyl cyanide-*p*-trifluoromethoxyphenylhydrazone for a final in-well concentration 0.5µM (“FCCP,” VWR Scientific), and a 100 µM mix of Rotenone (Sigma Aldrich) and Antimycin A (Alfa Aesar) for a final in- well concentration of 10µM of each (“R/A”). All port drugs were diluted to the desired concentration from 10 mM stock with DMSO and Seahorse medium and contained a final in-well concentration of 0.1% DMSO. Drug concentrations were determined using the manufacturer’s guidelines. OCR and ECAR measurements were completed at 4 stages; 1) with no injection, 2) after addition of oligo, 3) after addition of FCCP, 4) after addition of R/A. The effect of these drugs on mitochondrial function are described in **Figure 1**.

Because mtDNA CN varied by GWI status (described below), and mitochondrial content is likely to track mtDNA content, we normalized mitochondrial OCR measurement to 10˄6 mtDNA copies (mtDNA CN calculations described below). We mathematically combined OCR measurements to calculate the Bioenergetic Health Index [43], defined here as BHI = log((ATP-linked OCR x reserve capacity)/(proton leak x non-mitochondrial OCR)), and Respiratory Control Ratio [44], calculated as maximal (uncoupled) respiration divided by oligomycin-inhibited oxygen consumption.

### Isolation and quantification of WBC DNA

For mtDNA analyses, approximately 10 mLs were drawn into PAXgene Blood DNA tubes (Qiagen, 761115, containing 2mLs of a proprietary additive which prevents coagulation of the blood and preserves genomic DNA). PAXgene tube samples for mtDNA CN and damage assays were stored in a –20°C freezer for 24 hours, transferred to a -80°C ultra-freezer, shipped to Duke University on dry ice, and stored at -80°C until processing for DNA isolation. The frozen whole blood samples were thawed in a 37°C water bath for 15 minutes and then immediately processed. PAXgene Blood DNA kits (Qiagen, 761133) were used according to the manufacturer’s instructions to extract high molecular weight DNA. DNA yield and purity were initially analyzed using a NanoDrop ND-1000 (ThermoFisher), then further quantified using PicoGreen (ThermoFisher P7589) with a standard curve of a HindIII digest of lambda DNA (Invitrogen 15612-013) as described [54, 55].

This results in isolation of very high-quality, high-molecular weight total genomic DNA suitable for our DNA damage assay [56]. This isolated DNA was used for CN and damage assays described below.

### Measurement of mtDNA CN

We measured mtDNA CN using real-time PCR and plasmid-based standard curves for both mitochondrial and nuclear DNA counts, as we have described [56, 57]. All PCR reactions are conducted in duplicate or triplicate. To obtain mtDNA CN/cell, we divided the calculated mtDNA by 1,000: the real-time PCR reactions contained 2 μL of 3 ng/μL total DNA per reaction, for a total of 6 ng, and each diploid human cell contains ∼6pg total DNA. We also quantified nDNA CN, which we typically do as a quality check measure; the QPCR assay is carried with identical amounts of template (extracted) DNA, which is typically >99% nDNA because the nuclear genome is so much larger than the mitochondrial genome. However, in this study, we found a higher nDNA copy number per ng of total isolated DNA in veterans without GWI, as described in Results.

### Measurement of mtDNA damage

We measured mtDNA damage using a quantitative extra-long PCR-based assay that detects mtDNA by inhibition of DNA amplification [56]. This assay works by amplifying a very long (∼10 kb) region of the mitochondrial genome; any damage that inhibits the DNA polymerase used in the PCR reaction results in less amplification. Amplification of this long product is normalized to amplification of the small product (from the mtDNA CN assay) to account for any differences in numbers of mtDNAs. Then, any alterations in amplification of samples from veterans with GWI, compared to amplification of reference samples (in this case, from GVs who do not have GWI), are converted mathematically to a number of “lesions” (i.e., any damage that inhibits progress of the DNA polymerase) per 10 kb of mtDNA. Because this assay is PCR-based, it is specific for mtDNA damage without the need for differential isolation of mtDNA, which can result in artifactual results [58, 59].

### Statistical analyses

This is a non-randomized historical case-control study of GWI combining laboratory measured values of mitochondrial content and function with retrospective survey data pertaining to exposure and other risk factors related to GWI. Initial assessment of association of population characteristics, DNA measurements, or bioenergetic parameters with GWI status defined by the Kansas case definition were carried out by Chi-square tests, using the first value only for individuals who returned. Subsequently, we used a multivariate logistic regression with GWI status as the primary outcome to test the relationship with mitochondrial function, adjusting for important predictors. Older age is associated with mitochondrial change [60, 61], and may also reflect higher rank at time of deployment; higher rank in turn has been associated with better health among GVs [62, 63], and could potentially influence exposure as well. Therefore, age was evaluated in the relationship between GWI status and mitochondrial function. We used the average value for repeated mitochondrial function measurements as recommended by similar studies using a fixed outcome with few repeatedly measured predictors [64, 65]. We evaluated potential confounding variables (e.g., age, sex, body mass index, pyridostigmine bromide, etc.) prior to evaluating mitochondrial measures by examining univariate relationships by GWI status and considering covariates for a final model if their univariate relationship with GWI had a p-value under 0.2. Covariates where then evaluated for collinearity and then entered into a final model sequentially, with the most important covariates entering first. The final model was then evaluated for inclusion of each mitochondrial measurement. Potential outliers and influential observations were also evaluated by looking at deviance statistics.

## Results

We enrolled 121 veterans, and excluded seven due to potentially confounding health issues. Of the remaining 114, 80 met Kansas criteria for GWI, and 34 did not. 27 GVs returned for a second visit, and 26 for a third visit, allowing us to assess the consistency of measured parameters over time. In a small number of cases, specific analyses failed for some samples, resulting in smaller “n”s for specific assays. An overview of the characteristics of the participants is provided in **Table 1**. In general, younger veterans were more likely than older veterans to have GWI; strikingly, 0 of the 10 oldest veterans in our study had GWI. Veterans with GWI had a higher BMI and were more likely to smoke intensely, defined as daily smoking either at the time of interview or in the past. Finally, there was a strong association of recalled pesticide use, pyridostigmine bromide use, and exposure to chemical and biological warfare agents with GWI.

**Table 1.** Study population characteristics.

|  |  | All |  | GWI |  | non-GWI |  |  |
| --- | --- | --- | --- | --- | --- | --- | --- | --- |
|  |  | N | % | N | % | N | % |  |
| **Age** | 44-49 | 29 | 25.4 | 26 | 89.7 | 3 | 10.3 | +++ |
|  | 50-54 | 35 | 30.7 | 27 | 77.1 | 8 | 22.9 |  |
|  | 55-59 | 25 | 21.9 | 18 | 72.0 | 7 | 28.0 |  |
|  | 60-64 | 15 | 13.2 | 9 | 60.0 | 6 | 40.0 |  |
|  | 65+ | 10 | 8.8 | 0 | 0.0 | 10 | 100.0 |  |
| **BMI** | Normal (<25) | 11 | 9.7 | 4 | 36.4 | 7 | 63.6 | † |
|  | Overweight (25-29) | 51 | 44.7 | 36 | 70.6 | 15 | 29.4 |  |
|  | Obese (≥30) | 52 | 45.6 | 40 | 76.9 | 12 | 23.1 |  |
| **Gender** | Male | 102 | 89.5 | 71 | 69.6 | 31 | 30.4 |  |
|  | Female | 12 | 10.5 | 9 | 75.0 | 3 | 25.0 |  |
| **Pyridostigmine bromide** | None | 28 | 31.1 | 12 | 42.9 | 16 | 57.1 | †† |
|  | 1-6 days | 20 | 22.2 | 16 | 80.0 | 4 | 20.0 |  |
|  | 7+ days | 42 | 46.7 | 35 | 83.3 | 7 | 16.7 |  |
| **Pesticide Cream** | None | 40 | 43.5 | 21 | 52.5 | 19 | 47.5 | †† |
|  | 1-6 days | 14 | 15.2 | 10 | 71.4 | 4 | 28.6 |  |
|  | 7+ days | 38 | 41.3 | 33 | 86.8 | 5 | 13.2 |  |
| **Pesticide on Uniform** | None | 38 | 52.8 | 19 | 50.0 | 19 | 50.0 | †† |
|  | 1-6 days | 6 | 8.3 | 5 | 83.3 | 1 | 16.7 |  |
|  | 7+ days | 28 | 38.9 | 24 | 85.7 | 4 | 14.3 |  |
| **Smoking History** | None or not intense smoking | 78 | 68.4 | 50 | 64.1 | 28 | 35.9 | † |
|  | Intense smoking | 36 | 31.6 | 30 | 83.3 | 6 | 16.7 |  |
| **Alcohol Use** | None | 21 | 18.4 | 15 | 71.4 | 6 | 28.6 |  |
|  | 1-3 days/week | 81 | 71.1 | 60 | 74.1 | 21 | 25.9 |  |
|  | 4+ days/week | 12 | 10.5 | 5 | 41.7 | 7 | 58.3 |  |
| **IPAQ Score** | Q1 | 32 | 29.6 | 22 | 68.8 | 10 | 31.3 |  |
|  | Q2 | 26 | 24.1 | 21 | 80.8 | 5 | 19.2 |  |
|  | Q3 | 22 | 20.4 | 17 | 77.3 | 5 | 22.7 |  |
|  | Q4 | 28 | 25.9 | 15 | 53.6 | 13 | 46.4 |  |
| **Exposure to Chemical Warfare** | None | 35 | 55.6 | 14 | 40.0 | 21 | 60.0 | ††† |
|  | 1-6 days | 14 | 22.2 | 12 | 85.7 | 2 | 14.3 |  |
|  | 7+ days | 14 | 22.2 | 14 | 100.0 | 0 | 0.0 |  |

Note: Some variables have missing data due to non-response in the survey.
Chi-square tests to compare differences by GWI status: ††† p<0.0001; †† p<0.01; † p<0.05;

### DNA copy number

We previously reported increased mtDNA CN and mtDNA damage in veterans with GWI in a small study of 21 case and 7 control individuals [37]. Here, we replicated the finding of increased mtDNA (**Table 2**, **Fig. 2A**), when analyzing the participants on a binary (meeting Kansas criteria or not) case-control basis. mtDNA:nDNA ratio was approximately 9% higher in veterans with GWI than in veterans without GWI (**Table 2**, p = 0.08 by Chi square test). To test whether this was consistent across visits for individuals who returned, we carried out a repeated-measures ANOVA, and found that it was (p = 0.08). Interestingly, we found a higher nDNA copy number per ng of total isolated DNA in veterans without GWI. This result is unexpected, because under typical circumstances, most cellular DNA by mass is nuclear DNA. It is possible that the difference we observed may be explained by differences in platelet abundance in our samples: platelets are highly abundant, and do not contain nuclei (unlike other white blood cells), but do contain mitochondria and mtDNA. Therefore, high levels of platelets can significantly increase the proportion of mtDNA to total extracted DNA [66], and previous research has supported increased platelet count in veterans with GWI [67]. However, it is not clear that the *proportion* of platelets increases in GWI compared to other WBCs, as counts of many WBCs also increase in GWI [68]. Because of this uncertainty, we focus on mtDNA:nDNA ratios rather than absolute mtDNA/cell in our analyses of mtDNA amount, although we also report the latter (**Table 2**).

**Figure 2.**
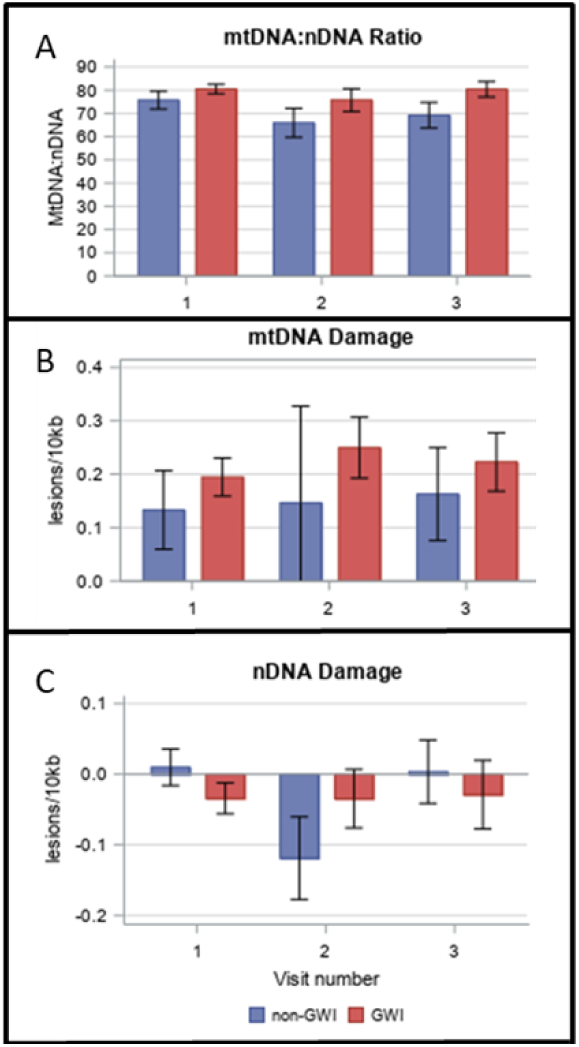
mtDNA copy number is increased in GWI, while mtDNA damage and nDNA damage are unchanged. A) mitochondrial and nuclear DNA CN from whole blood were measured with real­time PCR. B) mtDNA and C) nuclear damage were measured using a long- range quantitative PCR assay.

**Table 2.** Mean mitochondrial and bioenergetic values by GWI status.

|  | All | GWI | Non-GWI | p-value<br>(GWI vs non) |
| --- | --- | --- | --- | --- |
|  | Mean (95% CI) | Mean (95% CI) | Mean (95% CI) |  |
| mtDNA/6ng total DNA | 1049409 (1003055-1095764) | 1070078 (1013839-1126318) | 1003427 (919766-1087087) | 0.1906 |
| nDNA/6ng total DNA | 13527 (13311-13743) | 13400 (13132-13668) | 13766 (13400-14133) | 0.1119 |
| mtDNA:nDNA ratio | 78.2 (74.2-81.5) | 80.2 (76.0-84.3) | 73.6 (67.5-79.8) | 0.0824 |
| mtDNA Lesions/10 kb | 0.166 (0.11-0.22) | 0.183 (0.12-0.25) | 0.130 (0.03-0.23) | 0.3753 |
| nDNA Lesions/10kb | -0.029 (-0.059- -0.0003) | -0.035 (-0.072-0.0011) | -0.018 (-0.069-0.033) | 0.5791 |
| ATP-linked OCR/10<sup>6</sup> mtDNAs | 1.875 (1.60-2.15) | 1.619 (1.28-1.96) | 2.385 (1.92-2.85) | 0.0107 |
| Proton Leak /10<sup>6</sup> mtDNAs | 0.512 (0.42-0.61) | 0.457 (0.34-0.58) | 0.617 (0.45-0.78) | 0.1228 |
| Spare Respiratory Capacity/10<sup>6</sup> mtDNAs | 2.933 (2.21-3.66) | 2.570 (1.67-3.48) | 3.670 (2.37-4.98) | 0.1665 |
| Basal OCR/10<sup>6</sup> mtDNAs | 2.367 (2.01-2.73) | 2.075 (1.63-2.52) | 2.953 (2.33-3.57) | 0.0252 |
| Maximal OCR/10<sup>6</sup> mtDNAs | 2.933 (2.21-3.66) | 2.570 (1.67-3.48) | 3.670 (2.37-4.98) | 0.0856 |
| Non-mitochondrial OCR/10k cells | 1.396 (1.29-1.50) | 1.330 (1.20-1.47) | 1.514 (1.33-1.70) | 0.1151 |
| BHI | 0.992 (0.91-1.08) | 0.998 (0.89-1.11) | 0.982 (0.83-1.13) | 0.8575 |
| RCR | 3.880 (3.60-4.16) | 3.778 (3.43-4.13) | 4.067 (3.60-4.53) | 0.3214 |
| ECAR at baseline/10k cells | 3.021 (2.84-3.20) | 2.907 (2.68-3.13) | 3.267 (2.94-3.60) | 0.0397 |
| ECAR after oligomycin/10k cells | 3.469 (3.27-3.67) | 3.334 (3.09-3.57) | 3.755 (3.41-4.10) | 0.0741 |
| ECAR after FCCP/10k cells | 3.493 (3.35-3.64) | 3.405 (3.23-3.58) | 3.678 (3.42-3.93) | 0.082 |
| ECAR after RA/10k cells | 2.727 (2.59-2.87) | 2.623 (2.45-2.80) | 2.941 (2.69-3.19) | 0.0516 |

p-values based on Type 3 Chi-square tests to compare differences by GWI status Mitochondrial OCR measurements are normalized to mtDNA CN as described in Methods.

### DNA damage

Neither mtDNA damage nor nDNA damage was statistically different in veterans with GWI than in veterans without GWI (**Table 2**, **Figs. 2B and 2C**). This result was consistent across repeated visits (p = 0.38 and 0.58). For these analyses, no additional factors were considered in the analysis, and no individuals were considered statistical outliers, although a small number of individuals had mtDNA CN values that appeared anomalously low, increasing variance.

### Mitochondrial respiration

We quantified multiple measurements of mitochondrial oxygen consumption rate (OCR). These analyses provide a direct measurement of oxidative phosphorylation at baseline in resting cells, and also provide information on how much of this oxygen consumption can be attributed to production of ATP, how much is attributable to proton leak, how much OCR can be increased by mitochondrial uncoupling, and more (described in **Fig. 1**). We found significantly decreased oxygen consumption in white blood cells of veterans with GWI compared to veterans without GWI (**Fig. 3A**; results shown are corrected for mtDNA CN). Statistical analyses were carried out using a repeated measures analysis for basal, ATP-linked, maximal, spare, and proton leak-linked OCR. The full de-identified dataset is available in **Supplemental data file 1**. Basal OCR (p = 0.03) and ATP-linked OCR (p = 0.01) were statistically lower in veterans with GWI (**Figs. 3B and 3C)**. Maximal OCR (p = 0.09), spare respiratory capacity (p = 0.17), and proton leak (p = 0.12) were not statistically significantly different between the two groups (**Figs. 3D-F**). However, we note that differences in OCR after FCCP injection were much larger at the second and third readings after FCCP injection, and that OCR declined consistently after FCCP injection for the GWI population, but not the non-GWI population, suggesting that the maximal capacity and/or resilience to uncoupling challenge is in fact lower in GWI (**Fig. 3A**). BHI and RCR were not statistically different between the groups (p = 0.85 and 0.32, respectively). Various measures of OCR were relatively well-correlated with each other, likely explaining the lack of significant difference in the BHI and RCR indices. Overall, white blood cells from veterans with GWI used oxidative phosphorylation 30-40% less to make ATP than did white blood cells from veterans without GWI.

**Figure 3.**
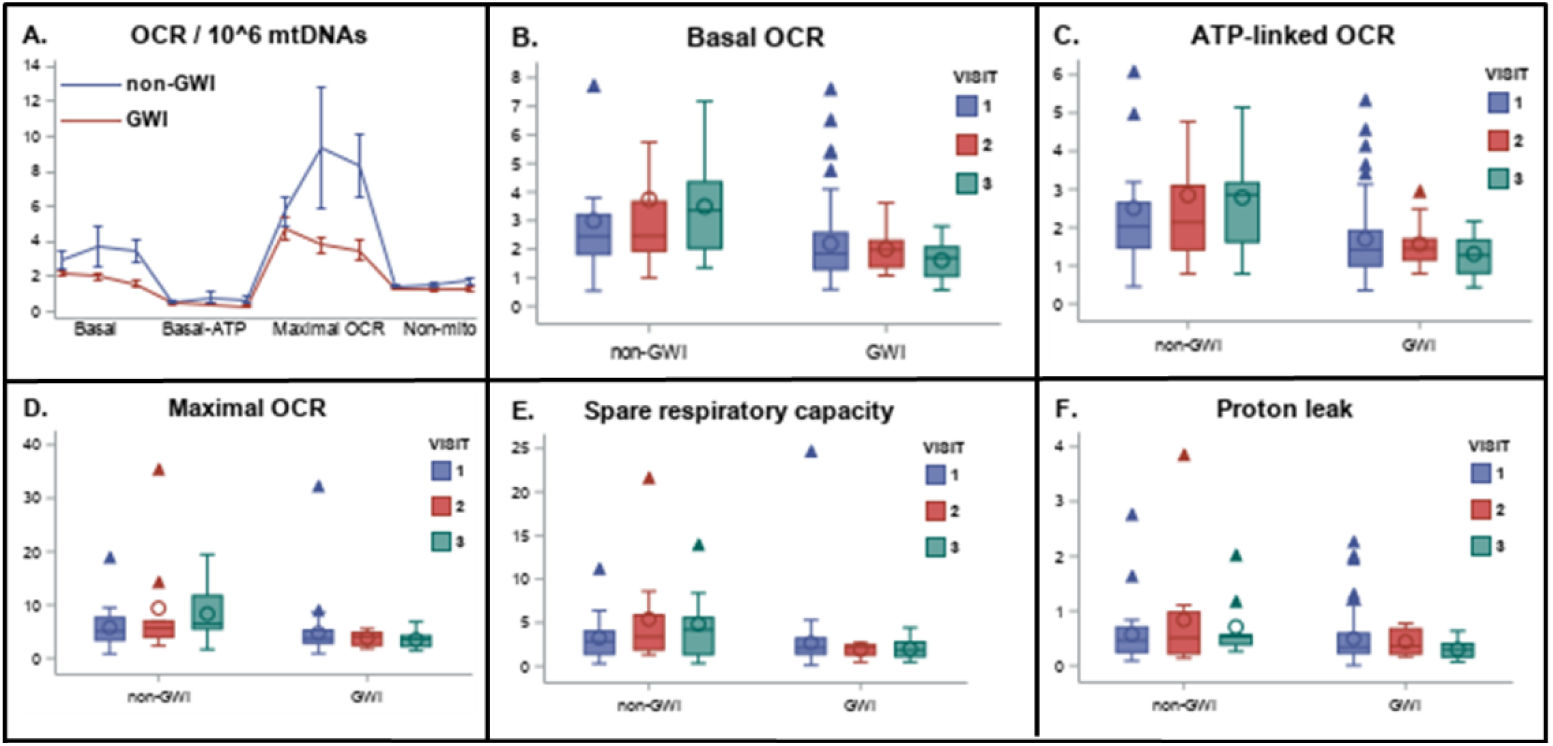
Decreased mitochondrial respiratory function in GWI. A) Average (with standard errors of the mean) oxygen consumption in all samples measured from veterans with and without GWI. Values are normalized to 1,000,000 mtDNA copies. The “basal” reading reflects the amount of oxygen being consumed by cells without the addition of any drugs. ATP-linked OCR is the amount of oxygen consumed upon injection of oligomycin, which inhibits ATP synthesis and therefore blocks oxygen consumption associated with converting ADP to ATP. Maximal respiration is respiration after injection of FCCP, which uncouples mitochondria. Non-mitochondrial respiration is the amount of respiration remaining after the electron transport chain is entirely inhibited with a combination of rotenone and antimycin A. See Figure 1 for a schematic illustration of the effect of the drugs. Spare respiratory capacity is the differences between maximal and basal; proton leak is the difference between ATP-linked and non-mitochondrial. Specific OCR-related parameters of particular interestare graphed by Kansas status and visit in panels B-F.

### Non-mitochondrial oxygen consumption

Non-mitochondrial OCR was unchanged in veterans with GWI (p = 0.11). This result is interesting largely in supporting the assumption that the mitochondrial effects we see are specific, as opposed to reflecting a generalized decrease in cellular function.

### Non-aerobic energy production

We also measured extracellular acidification (ECAR). ECAR serves as a proxy for glycolysis, an energy-producing process that does not require mitochondrial oxidative phosphorylation. ECAR is sometimes increased when oxidative phosphorylation is decreased, presumably as compensation. However, ECAR was either unchanged or reduced in veterans with GWI (**Table 2**). Therefore, the possibility that decreased respiration-based production of energy was offset by increased anaerobic energy production can be ruled out. ECAR measurements at baseline and after injection of various drugs were relatively well-correlated with each other.

### Repeatability of measurements over time, and variability in GWI vs non-GWI

We found that measurements of mtDNA CN, mtDNA:nDNA ratio, spare respiratory capacity and maximal OCR showed moderate replicability over time, with intraclass correlation coefficients of 0.43, 0.45, 0.57, and 0.47, respectively (ICC values were computed using the Shrout-Fleiss method [69]). On the other hand, proton leak, BHI, and RCR showed low ICC values: 0.01, 0.02, and 0.01, respectively. A second observation was that basal OCR, non-mitochondrial OCR, ATP-linked OCR, mtDNA damage and mtDNA:nDNA ratio all exhibited significantly higher ICC values for veterans without GWI compared to those with GWI, while ICC values were higher for nDNA damage, spare respiratory capacity, and maximal OCR for veterans with GWI compared to those without. ICC for mtDNA damage was approximately the same regardless of GWI status. Variability of some of these parameters over time is depicted in **Figure 4**, and ICC values are shown in **Figure 5**. **Figure 4** also depicts the higher variability in observed measurements for veterans without GWI, which is common across all measures (i.e., for every measure across all samples, not just those who returned for more than one visit, standard deviation was higher for veterans without GWI).

**Figure 4.**
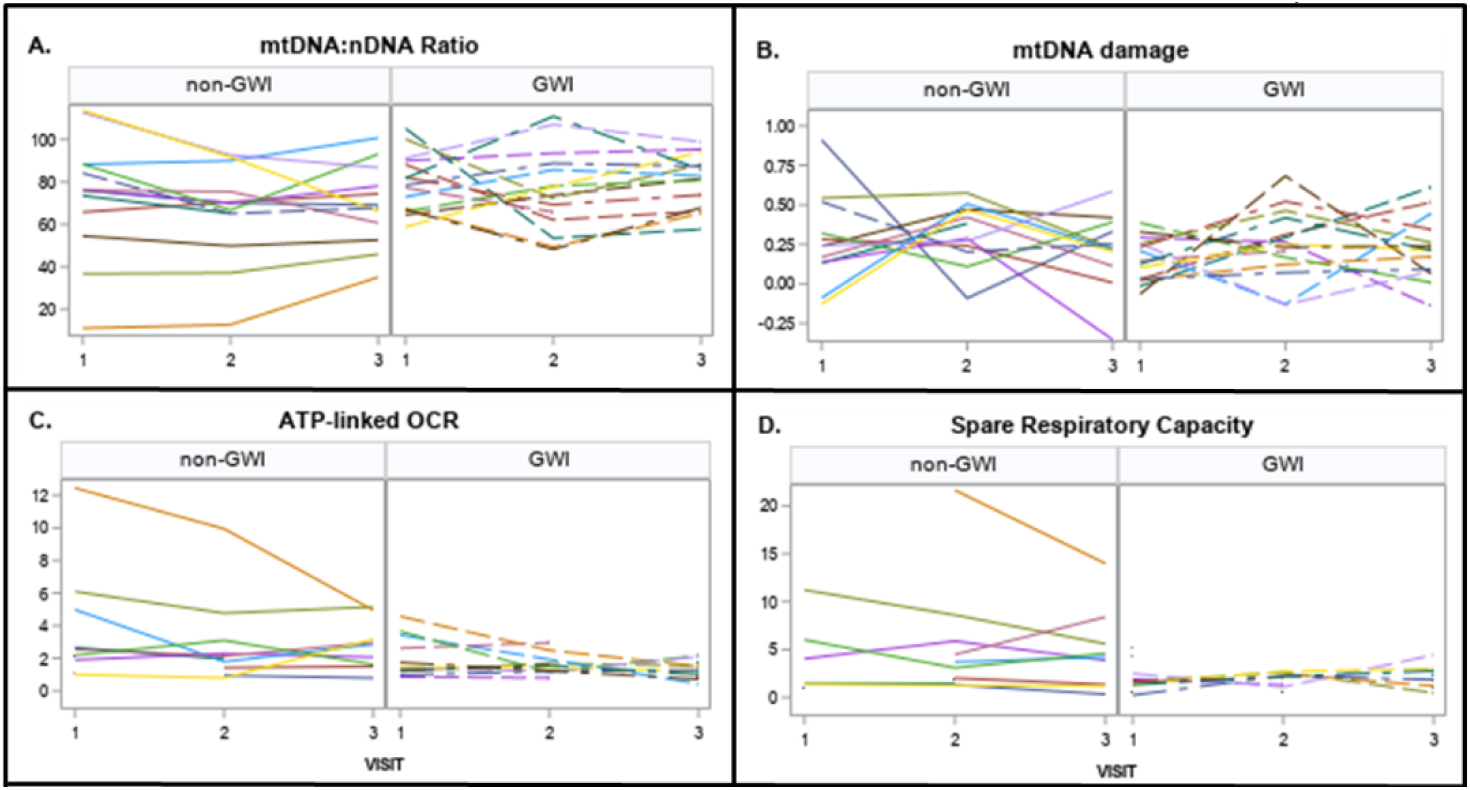
Repeatability of key mitochondrial parameters over time. Note that there is one outlier for l mtDNA damage that is not shown, to improve graphical readability; this subject is non-GWI with values **I** of -2.02, -1.94 and -0.44 for visits 1, 2 and 3 respectively.

**Figure 5.**
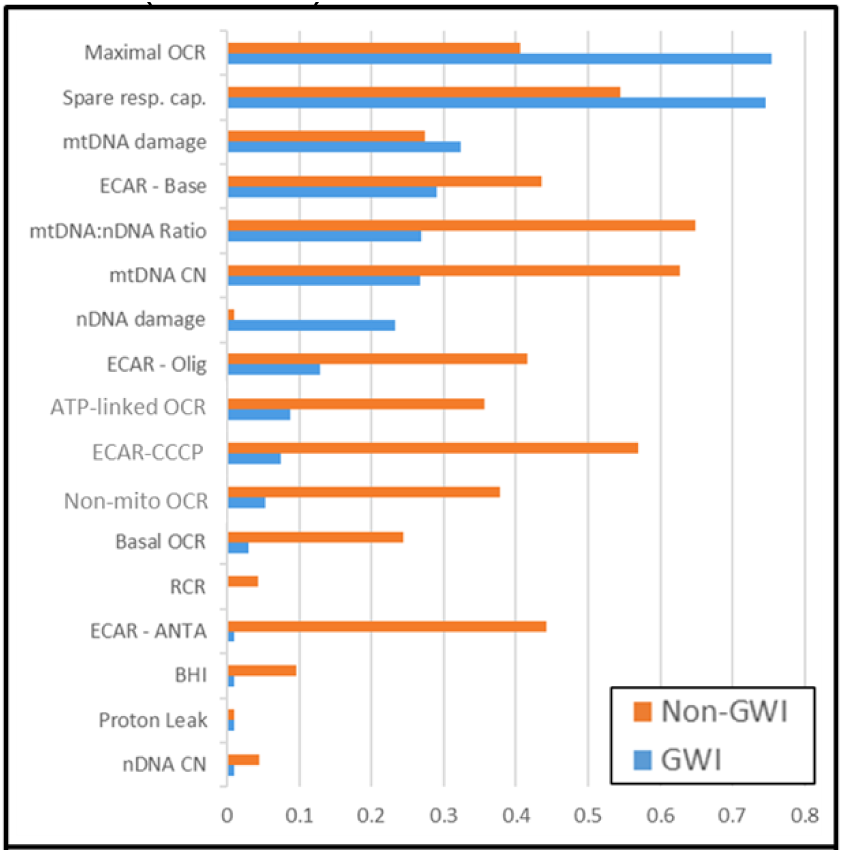
Intraclass Correlation Coefficients by GWI status. For graphing purposes, ICC estimates below zero were set to 0.01.

### Relationship of chemical exposures to mitochondrial parameters

To explore potential relationships between the exposures associated with GWI (**Table 1**) and mitochondrial parameters, we evaluated how key exposures were associated with mitochondrial parameters, controlling for GWI status using a repeated measures analysis. Higher recalled use of pyridostigmine bromide was associated with both decreased mtDNA CN (p=0.01) and mtDNA:nDNA ratio (p=0.05); use of pesticide skin cream was associated with higher mtDNA damage (p=0.08), while use of pesticide on uniforms was associated with lower nDNA damage (p=0.10); and exposure to chemical warfare was associated with increased non-mtDNA OCR (p=0.04). Predicted mitochondrial measures are depicted in **Supplemental Figure 1.** Chemical exposures were not associated with any other mitochondrial measure after controlling for GWI status.

### Effects of other variables on mitochondrial parameters

Mitochondrial function is modulated by factors other than chemical exposure (e.g., smoking, diet and exercise [70–74]). To reduce the potential for such factors to confound our results, as well as to inform future investigation of potential therapeutic interventions [75–77], we evaluated associations between mitochondrial parameters and age, sex, body mass index [BMI], international physical activity questionnaire index [IPAQ], smoking intensity, and alcohol consumption using repeated measures analysis. We found that heavier smoking was associated with higher mtDNA CN and higher mtDNA:nDNA ratio (p=0.02 in both cases) in models adjusted for GWI. To our surprise, we did not find a relationship between reported exercise levels or alcohol consumption and any of the mitochondrial parameters that we measured (p > 0.05). Of tested mitochondrial parameters, gender was only significantly related to mtDNA:nDNA ratio and BHI (females had higher values, p=0.045 and 0.036). Finally, after adjusting for GWI, age and BMI, no other factors were significantly related to mitochondrial parameters.

### Logistic regression analysis predicting GWI

To assess potential interactions between multiple factors in GWI, we carried out logistic regression analysis. Our multivariate model considered age, BMI, sex, smoking history, alcohol consumption, exercise (IPAQ), pyridostigmine bromide and chemical/biological exposures (use of pesticides on uniforms or on skin via creams, or exposure to chemical/biological warfare) as an initial base model. Model selection resulted in age and BMI as key covariates, along with pyridostigmine bromide use, pesticide exposure (on uniform or skin), and exposure to chemical/biological warfare; however, the latter three exhibited high collinearity and large differences in non-missing data. Thus, we fit four multivariate models (base model with age and BMI (n=114), base plus pesticide exposure (n=97), base plus pyridostigmine bromide (n=90), and base plus exposure to chemical warfare (n=63). Results from these models indicate higher odds of GWI for lower age, higher BMI, more exposure to pesticides and chemical warfare, and more use of pyridostigmine bromide (**Table 3**). The effect of DNA and bioenergetic parameters on GWI status was evaluated by including those measurements individually in each of the four models. DNA and bioenergetic parameters whose p-values were below 0.20 are shown at the bottom of **Table 3**. No mitochondrial or DNA measurement was significantly associated with GWI status at the 0.05 significance level after adjusting for age, BMI and exposure covariates. Higher levels of ECAR after rotenone + antimycin A, FCCP, and oligomycin injections were associated with lower odds of GWI in models that included pesticide exposure and exposure to chemical warfare. We note that no mitochondrial or ECAR measurement achieved significance in the model adjusting for age and BMI, which was the model that had the largest sample size evaluated.

**Table 3.** Multivariate Logistic Regression Model of GWI status and importance of mitochondrial measurements

|  | Model 1<br>(base) |  | Model 2<br>(base + Pesticide exposure) |  | Model 3<br>(base + pyridostigmine bromide) |  | Model 4<br>(base + chemical/biological agent exposure) |  |
| --- | --- | --- | --- | --- | --- | --- | --- | --- |
| Predictor | Log(OR) | ROC | Log(OR) | ROC | Log(OR) | ROC | Log(OR) | ROC |
| Intercept | 7.2729 <sup>††</sup> |  | 4.758 <sup>*</sup> |  | 3.2186 |  | 2.0007 |  |
| Age (years) | -0.1702 <sup>†††</sup> |  | -0.1164 <sup>††</sup> |  | -0.1105 <sup>†</sup> |  | -0.0915 <sup>*</sup> |  |
| Body Mass Index | 0.1010 <sup>**</sup> |  | 0.0838 |  | 0.1274 <sup>**</sup> |  | 0.1399 <sup>*</sup> |  |
| Pesticide on uniform or skin, REF=none |  |  |  |  |  |  |  |  |
| 1-6 days |  |  | -0.1906 <sup>**</sup> |  |  |  |  |  |
| 7+ days |  |  | 0.7913 <sup>**</sup> |  |  |  |  |  |
| Pyridostigmine bromide, REF= none |  |  |  |  |  |  |  |  |
| 1-6 days |  |  |  |  | 0.4969 <sup>*</sup> |  |  |  |
| 7+ days |  |  |  |  | 0.2943 <sup>*</sup> |  |  |  |
| Not exposed to chemical/biological agents |  |  |  |  |  |  | -1.1935 <sup>††</sup> |  |
|  | n=114 | 0.775 | n=97 | 0.795 | n=90 | 0.7807 | n=63 | 0.863 |
| Variables added to above models, one variable at a time |  |  |  |  |  |  |  |  |
| **DNA measures** |  |  |  |  |  |  |  |  |
| Nuclear DNA copies | -0.00012 | 0.7743 | -0.00025 | 0.796 | -0.00045 <sup>**</sup> | 0.8048 | -0.00043 <sup>*</sup> | 0.8772 |
| **Oxygen consumption measures** |  |  |  |  |  |  |  |  |
| ATP-linked/10<sup>6</sup> mtDNA copies | -0.4968 <sup>**</sup> | 0.8002 | -0.5187 <sup>*</sup> | 0.8052 | -0.3645 | 0.7993 | -0.3806 | 0.8714 |
| Non-mitochondrial/10K cells | -0.8013 <sup>**</sup> | 0.7933 | -0.5868 | 0.789 | -0.6502 <sup>*</sup> | 0.7902 | -1.3099 <sup>**</sup> | 0.8837 |
| Respiratory Control Ratio | -0.0562 | 0.7701 | -0.3334 <sup>*</sup> | 0.8036 | -0.1008 | 0.7748 | -0.4638 <sup>*</sup> | 0.8662 |
| Basal | -0.2928 <sup>*</sup> | 0.7902 | -0.3261 <sup>*</sup> | 0.8039 | -0.203 | 0.7941 | -0.1594 | 0.8597 |
| **Extracellular Acidification Rate** |  |  |  |  |  |  |  |  |
| Antimycin A | -0.878 <sup>**</sup> | 0.7634 | -1.1341 <sup>†</sup> | 0.7891 | -0.5226 | 0.7708 | -2.136 <sup>††</sup> | 0.8941 |
| Baseline | -0.5783 <sup>*</sup> | 0.7662 | -0.6286 <sup>*</sup> | 0.7928 | -0.3498 | 0.7822 | -0.8311 <sup>*</sup> | 0.8559 |
| CCCP | -0.7896 <sup>**</sup> | 0.7634 | -0.892 <sup>**</sup> | 0.7882 | -0.5403 | 0.7822 | -1.7846 <sup>†</sup> | 0.8837 |
| Oligomycin | -0.6073 <sup>**</sup> | 0.7655 | -0.7396 <sup>**</sup> | 0.7928 | -0.3591 | 0.7803 | -1.177 <sup>†</sup> | 0.8802 |

††† p<0.001; †† p<0.01; † p<0.05; \*\* p<0.10; \*; p<0.20

## Discussion

Our results indicate significantly decreased bioenergetic function in PBMCs from veterans with GWI. In particular, we measured decreased oxygen consumption associated with ATP generation as well as decreased extracellular acidification; both are consistent with less availability of energy. This was the case despite a small *increase* in mtDNA CN in GWI. Thus, our results are consistent with the hypothesis that cellular energetic insufficiency contributes to GWI. We also obtained strong support for a role for chemical exposure in GWI, as previously reported. Finally, we found that GWI was more common among younger veterans. Below, we discuss each of these and other points; we also discuss possible relevance of these results for diagnosis and treatment, limitations, and future directions for research.

### Decreased bioenergetic function in GWI

We report large decreases in essentially every measure of mitochondrial respiration in PBMCs from veterans with GWI. In particular, basal and ATP-linked oxygen consumption were lower, indicating that PBMCs in GVs with GWI are using oxidative phosphorylation less to produce ATP. Measures of spare and maximal respiratory capacity followed the same trend of being greatly decreased in GWI, suggesting less ability to increase oxidative phosphorylation when needed, but did not reach statistical significance. The steady decrease in OCR after FCCP injection in GWI, but not in non-GWI, is also consistent with mitochondrial function being less resilient to challenge in GWI. Respiratory control ratio and bioenergetic health index were not statistically decreased in veterans with GWI, likely because in each case, both the numerator and denominator are decreased in GWI. Measures of extracellular acidification, a proxy for glycolysis, an important non-mitochondrial source of energy, were also lower in GWI. Interestingly, some ECAR but no OCR measurements remained significant in the logistic regression model after adjusting for age and BMI; this may be driven by the fact that older veterans in our study were exclusively non-GWI, and had high OCR measurements. We also had very few GVs without GWI in our youngest age range. Future studies should consider age structure of the cohort in study design and recruitment. Our cohort was also unbalanced in terms of BMI, with a higher proportion of high-BMI veterans with than without GWI. The lack of significance of mitochondrial parameters after adjusting for age, BMI, and exposures will require additional study to fully understand, but in general, this could indicate either that mitochondrial function is in a causal pathway with one or more of these factors, or that they are simply well correlated.

Consideration of causal pathways suggests a number of interesting hypotheses. For example, if exposures causing GWI resulted in energetic deficiencies leading in turn to weight gain, accounting for BMI and exposures would also account for the changes in energetics in PBMCs.

Our finding of decreased respiratory function in PBMCs in GWI is consistent with previous literature suggesting mitochondrial dysfunction in GWI [35–38], as well as in rodent models [31, 33, 34]. They are also consistent with recent work in cell culture models. Delic *et al*. [78], using a neuronal cell culture model, found that combined exposure to N,N-diethyl-meta-toluamide (DEET), chlorpyrifos, and PB (a combination intended to mimic veterans’ exposure in the GW) caused a large decrease in mitochondrial spare respiratory capacity in 4 hours, at exposure levels that did not affect cell viability at either 4 or 8 hours. Yates *et al*. [79] differentiated induced pluripotent stem cells derived from veterans with or without GWI into forebrain glutamatergic neurons, and exposed them to diisopropylflurophosphate (an organophosphate pesticide used as a sarin mimic) and cortisol (to mimic stress); this cellular model of GWI reduced cellular energetics measured as produced reducing power and mitochondrial membrane potential.

Energetic deficiency is presumably worsened further by parallel decreases in non-mitochondrial energy production, measured by extracellular acidification. Although this decrease was smaller than the proportional decrease in OCR in GWI, and may be in part accounted for by decreased carbon dioxide production during oxidative phosphorylation [80], it is nonetheless important in indicating at the very least that glycolysis does not appear to be increased in a compensatory fashion, at least in PMBCs, as is often the case in mitochondrial disease. Of note, the earlier-mentioned study that identified increased blood lactate in GVs with GWI [35] found no difference in blood lactate levels at rest; blood lactate was only increased upon exercise. Unfortunately, we did not take post-exercise samples.

Taken together, our results are consistent with systemic cellular bioenergetic deficiency as a key aspect of GWI. This result is also consistent with important symptoms observed in GWI, such as fatigability [4, 81, 82]. If the reduced OCR we observed in PBMCs is also present in muscle cells, for example, this may translate to less tolerance of energetic stress, resulting in exercise-induced fatigue or post- exertional malaise reported in GWI. Although our measurements were for practical reasons made in blood samples, not in tissues associated with GWI symptoms (muscle cells, nerves, etc.), our results are an important addition to previous literature because they directly demonstrate significant decreases in the principal function of mitochondria, energy generation.

### Increased mtDNA content in GWI

The increase we observed in mtDNA content is consistent with results from Yates *et al*. [79], who using the cell culture model described above found evidence for increased mitocondrial number in cells from veterans with GWI and cells challenged with GWI-relevant stressors. The increase in mtDNA CN could result at least in part from a compensatory program that increases mtDNA biogenesis (i.e., production of mitochondria, including mtDNA). mtDNA may be degraded upon damage [83], which could lead to a decrease in mtDNA CN and potentially in lower production of the mtDNA-encoded components of the electron transport chain. However, as we and others have discussed in some detail [45, 46], cellular stress often results in an increase in mitochondrial biogenesis (i.e., production of mitochondria, including mtDNA), such that an increase in mtDNA CN is a common occurrence in response to cellular stress and mitochondrial dysfunction. This presumably serves to compensate for decreased mitochondrial functionality, as appears to be the case in our cohort, by increasing mitochondrial capacity and/or replacing degraded components. Other factors that could contribute to the increase in mtDNA CN that we observed are an increase in platelet count as a proportion of WBCs in GWI, an increase in cell-free mtDNA in blood, or a decrease in lysosomal degradation of mitochondria. There is evidence for decreased lysosomal activity in brains of mice exposed to a GWI-mimicking mixture of chemicals [84]. Sepsis can result in release of mtDNA to blood [85], as can mitochondrial disease [86], and we speculate that milder inflammatory conditions such as those observed in GWI [87] may also result in systemic release of mtDNA, which may further exacerbate inappropriate immune response [86, 88, 89]. High levels of platelets can also significantly increase the proportion of mtDNA in total extracted DNA since they do not contain nuclei, and previous research has supported increased platelet count in veterans with GWI [67]. However, it is not clear that the proportion of platelets increases compared to other WBCs, as counts of many WBCs also increase in GWI [68]. We did not assess measures of mitochondrial biogenesis, platelet count, cell-free mtDNA, or lysosomal activity in WBCs in this study.

### Absence of detectable mtDNA damage in GWI

In a previous, smaller study, we reported mtDNA damage in circulating PBMCs in GWI [37]. This observation was not supported in this larger study. We also did not detect increased nuclear DNA damage.

### Variability in mtDNA CN and damage measurements

Because mtDNA CN and damage are increasingly being employed as biomarkers, it is important to consider sources of variability. We had fair consistency in some measurements made in the same individuals over time, which is encouraging. However, we also saw significant interindividual variability in these measures (interestingly, with greater interindividual variability in individuals without than with GWI). We similarly saw high interindividual variability in previous studies [55, 90]. That this large interindividual variation is biologically real as opposed to technical is supported by two lines of evidence. First, we previously saw a striking difference in averages between three different communities in Peru, despite high variability within each community [55], indicating that a significant portion of the variability was non-random. Second, in the current study, we identified two individuals with a surprisingly low mtDNA CN values. One of these individuals met Kansas criteria for GWI, and had 20% of average mtDNA CN; the other did not have GWI and on three separate visits had 11%, 13%, and 48% of average mtDNA CN. The individual who returned twice is particularly informative; the fact that all three reads were far below average indicates that it is highly unlikely that a technical error resulted in the same large deviation from the mean in all three analyses (DNA isolations and PCR reactions were carried out at different times for the different visits). Collectively, we suggest that these results indicate that important contributors to mtDNA CN and damage (and sources of variability in these parameters) remain unexplained.

### Exposures that may contribute to GWI

It is unfortunately challenging to understand which exposures might have caused GWI. Exposures were not carefully tracked or measured at the time, and the chemicals to which veterans are known to have been exposed are not long-lived in the body. Previous work has suggested a role for exposure to PB and several pesticides [91, 92], and our results are supportive of those reports. Steele et al. [93] found a stronger association between PB use and GWI in individuals with butyrylcholinesterase gene variants that may confer less ability to inactivate organophosphate and carbamate pesticides. There is also evidence from mouse studies that genetic differences may affect susceptibility to the development of GWI as modeled by co-administration of corticosterone and DPF [94]. Finally, a recent study also makes a strong case for exposure to low levels of nerve gas playing a role in GWI [95]. Although that was a correlative study, the mechanistic plausibility of the association is greatly strengthened by the fact that individuals with a lower-activity genetic variant of in the paraoxonase 1 gene (encoding a protein that is rate-limiting for metabolizing many nerve agents) had a ∼2-fold higher risk of GWI if they recalled hearing alarms indicating possible nerve gas exposure. We asked if veterans had exposure to chemical or biological warfare agents, but did not ask specifically about hearing chemical alarms, which would have permitted a closer comparison to this recent study. Nonetheless, we did find a strong association between recalled exposure to chemical or biological warfare agents and GWI, consistent broadly with the results of that study. Even after adjusting for age and BMI, exposure to chemicals (pesticides, PB or chemical warfare) increased the risk for GWI and significantly improves the sensitivity and specificity of predicting GWI, as indicated by the receiver operating characteristic curve (ROC) increasing from 0.77 to 0.8, 0.78 and 0.86, respectively. Finally, the strength of the associations we observed with PB, pesticide, and chemical warfare agent exposure might have improved if we had carried out genetic analysis.

### Other demographic factors

We were surprised to find that GWI was more prevalent among younger veterans in this study, given that mitochondrial function decreases with age (discussed below). We speculate that this may be a result of younger veterans being more likely to be deployed, as previously reported [96]. We were also surprised to find no effect of exercise on mitochondrial function (after controlling for GWI); however, while there is long-standing evidence that exercise affects mitochondrial function [97], we found only one small study specifically addressing effects of exercise on mitochondrial respiration in WBCs [98]. Finally, we note that other factors, such as psychological state [99, 100], may affect mitochondrial function.

### Potential for bioenergetic biomarkers in GWI diagnosis and therapy

Overall, the improved understanding of the degree to which mitochondria are altered in GWI provided by these results supports the likelihood that chemical exposures or other stressors that are capable of affecting mitochondria contributed to the development of GWI. Ideally, such measurements could also contribute to diagnosis, especially given the difficulties experienced by some veterans in being diagnosed with GWI in order to access appropriate care. The search for good biological biomarkers that can be used diagnostically for GWI is ongoing [101]. A biomarker is defined by the National Academy of Sciences as a “xenobiotically-induced variation in cellular or biochemical components or processes, structures, or function that is measurable in a biological system or sample” [102]. A good biomarker is sensitive, specific, reproducible, feasible to measure, and clearly related to health outcomes. Based on these criteria, we do not think that mtDNA CN, OCR, or ECAR could serve as stand-alone, completely unique-to-GWI biomarkers, at least without additional technical improvements. Mitochondrial parameters are altered in other conditions as well, limiting specificity. Indeed, mitochondrial parameters have been described as “canaries in the coalmine” [9, 25, 43, 103, 104], for this reason. We are aware of no previous studies evaluating ECAR as a biomarker, but note that in our studies, ECAR showed more promise than OCR in that statistically significant associations with GWI were maintained for some measurements even after adjustment for age and BMI. A second limitation is sensitivity: the change in mtDNA CN is relatively small, and the changes in OCR measurements in particular, while large, are variable among individuals. However, because they are quantifiable, objective, and mechanistically relevant, we do think that they could be a powerful addition to other markers and criteria used to diagnose GWI or assess improvement upon treatment. Because we found that some of these markers are more variable over time in GWI, making regular measurements will likely be important.

The urgency of understanding mitochondrial dysfunction in GWI is heightened by the age of these veterans. Unfortunately, mitochondrial function declines in old age in any case. A recent paper showed very dramatic decrease in mitochondrial respiration in PBMCs from adults aged 65-89 vs 53-64 [61]. In fact, mitochondrial dysfunction is a leading candidate for the aging process in general, and is associated with many age-related degenerative diseases [105–107], suggesting that veterans with GWI will experience an exacerbation of normal aging-related mitochondrial decline [108]. Furthermore, there is some evidence that GVs are aging at an accelerated rate [109]. This will exacerbate both symptoms, and the already-increasing difficulty of diagnosing GWI in aging veterans. Aging-related decline in mitochondrial function may be further exacerbated by physical inactivity. Exercise is used as therapy for many mitochondrial diseases [110, 111], and there is consensus from the Mitochondrial Medicine Society for exercise as a primary treatment [112]. Research into treatments for mitochondrial diseases is an active area of research [75, 76], and a significant number of intervention treatments are currently being tested for GWI, including exercise [113]. As reported in the GWI Landscape document [114], many of these are aimed at protecting mitochondrial function or reducing oxidative stress, derived primarily from mitochondria. However, most outcome measurements in these studies are self-reported function. Such studies could benefit from the ability to assess how objectively-measured mitochondrial function responds to treatment. While we observed significant variations in mitochondrial parameters between individuals, we also observed that some measurements were relatively steady over time in the same individuals, suggesting that mitochondrial function in PBMCs could serve as an objective readout of efficacy of treatment, especially given the relatively rapid turnover of most white blood cells.

Notably, GWI presents with a complex mixture of symptoms (not unlike many other chronic diseases and conditions), and evidence exists for molecular and cellular mechanisms other than mitochondrial dysfunction, including immune dysfunction, neuroinflammation, and more. While some of these mechanisms likely also involve mitochondria (e.g., mitochondria play critical roles in the immune response and in neuronal function), we recognize that etiology may include additional, non- mitochondrial mechanisms, at least in some veterans. If this is the case, therapies may also need to be personalized (as, again, is common in other complex diseases). An objective, quantifiable marker of mitochondrial (dys)function should assist in identifying those veterans who are most likely to respond to a mitochondrial-targeted therapy.

### Limitations and future directions

As mentioned above, a limitation associated with our results is that it they were obtained in circulating white blood cells, as opposed to cells from tissues most often affected in GWI. Relatedly, we used a pooled mixture of WBC types, which requires less processing than analysis of individual cell types. However, this means that our results may be diluted or skewed by changes in specific WBC populations (monocytes, lymphocytes, and neutrophils, as well as platelets, although platelets were excluded from our OCR analyses). The rationale for examining sub-populations is that these cell types differ in terms of circulating half-life, amount present in blood, mtDNA CN, and mitochondrial respiration. For example, neutrophils are highly glycolytic, while platelets and monocytes rely heavily on oxidative phosphorylation [115]. Based on such basic cell type differences, there are two fundamental reasons that separation of cell type would be beneficial. First, the proportion of each type of WBC is not constant person to person; thus, since metabolism and mtDNA CN vary by cell type, changes in WBC composition could create a misleading appearance of altered mitochondrial function. Second, differences associated with GWI may occur only in some cell populations. For example, because neutrophils are highly glycolytic to begin with, they are unlikely to show any decrease in mitochondrial respiration in GWI, and including them in a total WBC mitochondrial respiration measurement will dilute any differences that occur in other WBC types. Thus, valuable future directions would be analysis of mitochondrial status and function in specific cell types, and ideally in affected tissues. Another interesting possibility is analysis of cell-free mtDNA in blood. Recent evidence indicates that cell-free mtDNA may be released as a result of cellular and mitochondrial stress [116–118], and therefore could potentially serve as a biomarker of such stress. Another interesting future direction would be to assess whether mitochondrial dysfunction is more strongly associated with some GWI-related conditions than others (e.g., mitochondrial dysfunction may be more causally linked to neurological and fatigue domains, than skin-related domains).

A striking aspect of GWI is its persistence in veterans decades after the end of the conflict. A related and perplexing question is why mitochondrial function persistently altered in GWI. None of the chemicals for which there is a strong association with GWI are highly persistent in the human body, and other environmental factors presumably also changed when deployments were completed. It may be that once established, by mitochondrial toxicity or other mechanisms, mitochondrial dysfunction is simply a defining feature of GWI. It is possible that gene regulatory pathways that regulate the biological processes altered in GWI (inflammation, mitochondrial function, etc.) were reset during the GW. There is strong evidence from various animal model systems for mitochondrial stressors resulting in long-term alterations in mitochondrial function [119–122]; an important future direction would be to interrogate mechanisms such as epigenetic patterning [123] that could mediate persistently altered mitochondrial function.

### Conclusions

Our results demonstrate significantly decreased bioenergetic function in GWI. Strengths of the study include comprehensive analysis of bioenergetic function and incorporation of normalization of mtDNA copy number in a relatively large cohort of GVs. In combination with previous reports of altered mitochondrial markers in both blood and affected tissues in veterans with GWI, as well as in cell culture and animal models of GWI, our results are supportive of systemically altered mitochondrial function in GWI. Together with an absence of any sign of increase in glycolysis, these results suggest that at least some of the symptoms that characterize GWI are based on cellular energetics deficiency. We hope that these results will support current diagnosis of GWI, and that they will improve the likelihood and effectiveness of treatment of current veterans. The variability in measurements between individuals with GWI limits the use of these measurements as stand-alone biomarkers of GWI *per se* without further technical improvements. However, we propose that they could serve as one additional criterion, and potentially one that could be used to objectively assess progression or the effect of interventions given relatively good consistency in some measurements over time. In the context of therapy, our results suggest that it would be worthwhile to test treatments aimed at mitochondrial function, potentially including both drugs and lifestyle interventions (e.g., exercise) being tested for mitochondrial diseases. Finally, we hope that our results will be informative with regard to reducing potentially deleterious chemical exposures in the future, and to the extent that such exposures are unavoidable, minimize them and support treatment for any resulting toxicity.

## Acknowledgements

We gratefully acknowledge the Veterans who volunteered to participate in this study. We would also like to thank the research team that helped recruit volunteers and acquire data including Nancy Eager, Matthew Watson, Wei Qian, and Helene Domanski. The authors would also like to thank the Veterans Affairs (VA) Cooperative Studies Program 585 Gulf War Era Cohort and Biorepository (GWECB CSP#585) for use of portions of their questionnaire. This work was supported by the Office of the Assistant Secretary of Defense for Health Affairs through the Gulf War Illness Research Program under Award No. W81XWH-16-1-0663, and supported in part by the Merit Review Award # I01 CX001329 from the United States Department of Veterans Affairs Clinical Sciences Research and Development Service.

The contents including the opinions, interpretations, conclusions, and recommendations, are those of the authors and are not necessarily endorsed by the Department of Defense, U.S. Department of Veterans Affairs or United States Government.

---

**Supplementary Figure 1.**
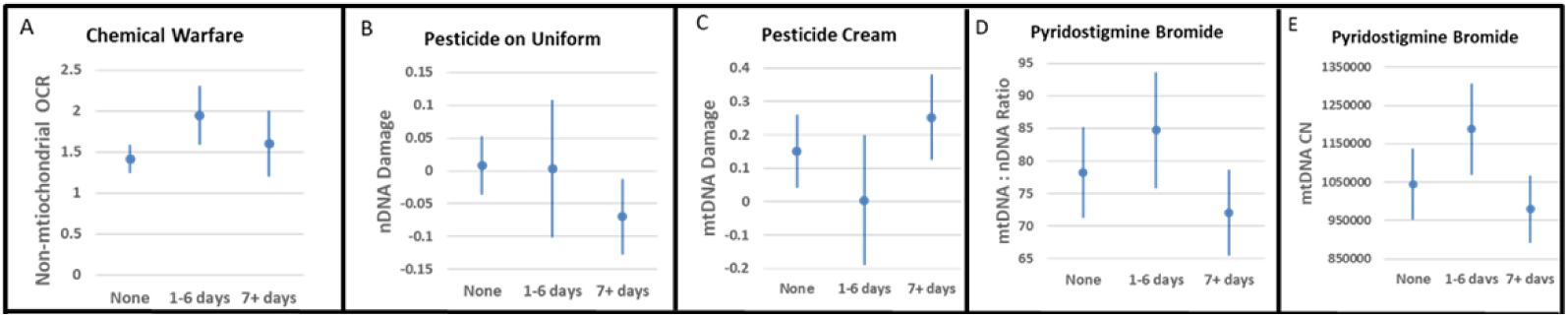
Exposures that were associated with mitochondrial parameters after controlling for. GWI status. Mean effects and 95% confidence intervals for days of chemical exposures, adjusting for GWI status.

